# Statistical evidence for common ancestry: New tests of universal ancestry

**DOI:** 10.1101/036327

**Authors:** Bret Larget, Cécile Ané, Martin Bontrager, Steve Hunter, Noah Stenz, David A. Baum

## Abstract

While there is no doubt among evolutionary biologists that all living species, or merely all living species within a particular group (e.g., animals), share descent from a common ancestor, formal statistical methods for evaluating common ancestry from aligned DNA sequence data have received criticism. One primary criticism is that prior methods take sequence similarity as evidence for common ancestry while ignoring other potential biological causes of similarity, such as functional constraints. We present a new statistical framework to test separate ancestry versus common ancestry that avoids this pitfall. We illustrate the efficacy of our approach using a recently published large molecular alignment to examine common ancestry of all primates (including humans).

We find overwhelming evidence against separate ancestry and in favor of common ancestry for orders and families of primates. We also find overwhelming evidence that humans share a common ancestor with other primate species.

The novel statistical methods presented here provide formal means to test separate ancestry versus common ancestry from aligned DNA sequence data while accounting for functional constraints that limit nucleotide base usage on a site-by-site basis.

---

The notion of common ancestry (CA) among all species dates most prominently to the work of Charles Darwin in the foundational text *The Origin of Species* (Darwin 1859). More than 150 years later, the primary ideas about evolution put forth by Darwin are supported by voluminous qualitative evidence (Coyne 2009), which explains why biologists almost universally accept CA and other tenets of evolutionary theory as well-supported scientific truths. Nevertheless, CA, especially with regard to the relationship of humans to other species, remains controversial among a large fraction of the general public. Perhaps motivated in part by this controversy, some researchers have sought to develop formal statistical arguments to test the hypothesis of CA.

The first such statistical test of CA (*Penny et al.* 1982), published in 1982, used as evidence the highly unlikely topological agreement among the most-parsimonious trees for five separate proteins sampled from the same 11 taxa, assuming that all possible rooted trees among 11 taxa are equally likely and independent under a null hypothesis of separate ancestry (SA). This test looks for tree structure and is not obviously appropriate for assessing universal CA — the contention that all tips are connected through evolutionary descent rather than originating from two or more separate origins. A discussion of appropriate principles by which to test CA statistically left the question unresolved (Sober and Steel 2002; *Penny et al.* 2003). To fill this gap, two recent papers have developed formal statistical arguments for CA (Theobald 2010a; W. Timothy J. White 2013). In the first of these papers, Theobald applied likelihood ratio tests (LRT), Bayes factors, and the Akaike information criterion (AIC) to the three domains of present-day life (Archaea, Bacteria, and Eukarya) and found overwhelming evidence for universal CA. While not questioning his conclusions, several papers subsequently raised objections to Theobald's statistical methods (Yonezawa and Hasegawa 2010; Eugene V. Koonin 2010; Yonezawa and Hasegawa 2012; de Oliveira Martins and Posada 2014). The most compelling among these objections was that the results of the tests are a trivial consequence of significant similarity among the sequences.

There are at least two ways in which sequence similarity can mislead formal statistical tests and overstate the strength of evidence in favor of CA. First, the process of sequence alignment injects gaps into the sequences to account for an unknown history of insertions and deletions along different lineages. The gaps are selected so that the aligned sequences are maximally similar subject to some constraints. If the alignment process mistakenly separates bases in different species that should be directly compared, then the strength of evidence for CA could be overstated. However, this possible source of potential overconfidence is negligible when there is little alignment uncertainty and is easily handled by excluding regions where alignment uncertainty is relatively high. A second and more serious concern is if sequence similarity were due to a biological cause other than common ancestry. Yonezawa and Hasagawa demonstrated that Theobald's methods favored CA over SA even when comparing unrelated mitochondrial genes (*cytb* and *nd2* from cow, deer, and hippopotomous) that were aligned without gaps by simply truncating the end of the longer sequence (Yonezawa and Hasegawa 2010). In this instance, Theobald's methods were misled by the distinct preferences in nucleotide base composition by codon position common to all protein-coding mitochondrial genes (Theobald 2010b). In particular, nonidentical base distributions at each site contradict a key assumption underlying the likelihood model used for SA by Theobald. One can also imagine when considering a hypothesis of SA that comparable genes with similar functions in separate species could have similar sequences due to functional constraints rather than to CA. In light of all of this criticism, the community remains without a thoroughly convincing statistical method to demonstrate universal CA, whether among all domains of life or for more specific sets of species.

To address this void, we propose novel statistical tests for CA that are related to the approaches put forth by Theobald, but avoid objections raised by others. In particular, using our methods, an exact agreement among sequences is uninformative and unable to reject SA because complete agreement is as equally well explained by strong functional constraints as by CA. The first method we propose uses the Bayesian information criterion (BIC) under a novel likelihood model that allows for site-specific stationary distributions. For the second method, we develop a bootstrap approach to simulate the null distribution of a parsimony-based test statistic to quantify statistical evidence.

Rather than addressing the question of a single universal common ancestor for all domains of life, we decided to focus instead on primate CA. This choice facilitates method development because of the ready availability of large aligned data sets. Additionally, it becomes reasonable to ask the specific question of how strongly molecular sequence data support the inference that the human species shares CA with other primates. The methods in this paper are separate and distinct from those in companion papers (*Baum et al.* 2015; *Bontrager et al.* 2015) that examine CA in primates using alternative statistical methods. Before describing our methodology in more detail, we look a bit more closely at the Theobald approach to clarify how we overcome the objections raised by previous authors.

## Likelihood in Theobald (2010)

Theobald finds support for CA using LRTs, Bayes factors, and AIC. Each of these tests depends on the evaluation of the likelihood of the sequence data under a model. To illustrate the use of likelihood in this setting, consider the following 10-base sequence which is found at the beginning of both the human and chimpanzee *ABCA1* gene: TGTGTCTCAC. Using maximum likelihood, the stationary base composition would be estimated as (0.1, 0.3, 0.2, 0.4) for bases A, C, G, and T, respectively, for both the CA and SA models. For CA, this distribution is estimated by the observed relative frequency of nucleotide base usage over the entire data set. For SA, the human and chimp distributions are estimated separately. However, because, in this case, the nucleotide usage is identical for both sequences, the estimated distributions also agree. Under the CA model, the observed data would have the following maximum likelihood

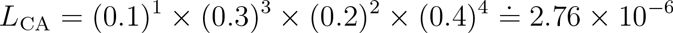

where the branch length and susbstitution parameters would be estimated so that there is probability one of no sequence changes from common ancestor to both chimpanzee and human. Under the SA model, this ancestral sequence would be generated at random twice and the maximum likelihood would be

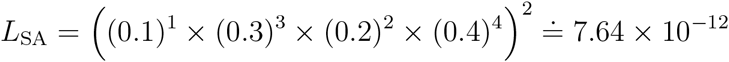

resulting in a difference in log likelihoods of 12.8, with the CA model more likely, even though the SA model has more parameters than the CA model (potentially two distinct stationary distributions and potentially two distinct models for nucleotide substitution). Neither model is nested within the other and conventional distributional results about LRT statistics do not apply. The reason the models are not nested is that no parameters of the larger SA model guarantee a common ancestral sequence for both humans and chimpanzees. Nevertheless, any of Theobald's three approaches would have found these short sequences to be statistically significant in favor of CA versus SA on the basis of the sufficiently large difference in log likelihoods. Theobald finds strong evidence for CA because comparable sequences are more similar to each other than would be expected if they were generated independently using the same distribution for each site in the sequence.

We make a fundamentally different assumption about the nature of ancestral sequences. Under the SA hypothesis, we suppose that the collection of observed sequences are shaped by unspecified biological constraints. Specifically, we assume that each site in an ancestral sequence has its own probability distribution of possible bases which we estimate independently of data from other sites.

In this small example with the first ten bases of the *ABCA1* gene for humans and chimps, there is no sequence variation. With our approach, the SA model requires the exact sequence TGTGTCTCAC at the two ancestral sequences and the maximum likelihood fitted model allows no nucleotide substitution. The null sampling distribution of human and chimpanzee sequences under this estimated model has no changes among the sequences with probability one and the observed data is consistent with the null hypothesis of SA.

Essentially, our approach completely discounts all sites that have no variation, thus avoiding the primary criticisms of Theobald's methods. We derive information for CA versus SA from sites that vary. Our first method uses BIC with a likelihood model with site-specific stationary distributions. Our second method simulates the null sampling distribution of a parsimony-based measure of sequence differences among groups assumed to have SA generated from a site-specific model of SA and compares this distribution to the observed parsimony measure from the original sequences.

## Primate phylogeny

A recent publication (*Perelman et al.* 2011) contains a molecular phylogeny of primates created using 54 nuclear genes and 191 taxa including 186 primate taxa from an alignment of 34,941 base pairs that the authors reduced from a larger alignment after discarding sites with great alignment uncertainty. Sequence data included roughly equal amounts of coding and noncoding sites, mostly from autosomal regions of the genomes, but with a few thousand sites from both X and Y chromosomes. No taxon was sampled for all 54 genes (humans are the most sampled with 53 genes) and many taxa have long stretches of missing data. We use the same alignment of data subject to further modifications to eliminate outgroups and combine subspecies as described in Methods, resulting in a final alignment with 178 taxa, each of which we consider to be a single species, and 34,860 sites.

The order Primates is divided into two suborders, Haplorrhini (dry-nosed primates, including tarsiers, New World and Old World monkeys, and apes) and Strepsirrhini (wet-nosed primates, including galagos, pottos, lorises, aye-aye, and lemurs). The primate taxonomy literature does not agree uniformly on the definitions of families. We choose to follow the most widely held conventional taxonomy of the 16 families described in Table 1 as our working definition. Switching to alternative family definitions as used elsewhere in the literature (Groves 1991; Hill 1966; *Perelman et al.* 2011) would not alter our primary results about CA.

**Table 1:** **Primate families in the suborders Haplorrhini and Strepsirrhini.** The numbers of genera and species correspond to the counts among the 178 species in the data set used in this study. There are additional species and genera among living primates that are absent from this data set.

| Suborder | Family | # of genera | # of species |
| --- | --- | --- | --- |
| Haplorrhini | Cercopithecidae | 21 | 62 |
|  | Hylobatidae | 3 | 8 |
|  | Hominidae | 4 | 6 |
|  | Callitrichidae | 6 | 20 |
|  | Aotidae | 1 | 4 |
|  | Cebidae | 2 | 10 |
|  | Atelidae | 4 | 14 |
|  | Pitheciidae | 4 | 14 |
|  | Tarsiidae | 1 | 2 |
| Strepsirrhini | Lepilemuridae | 1 | 5 |
|  | Cheirogaleidae | 3 | 3 |
|  | Indriidae | 2 | 5 |
|  | Lemuridae | 4 | 13 |
|  | Daubentoniidae | 1 | 1 |
|  | Lorisidae | 4 | 6 |
|  | Galagidae | 2 | 5 |

## BIC test for common ancestry

Both formal testing methods we introduce can be used to compare CA versus SA at multiple levels of taxonomy. For each such test, we partition the taxa into two or more groups. Both hypotheses assume a tree-like speciation process within groups with species within each group descended from a most recent common ancestor. The difference is that universal CA assumes that the groups are all related in a single tree, whereas SA posits separate origins for each group (see Figure 1).

**Figure 1:**
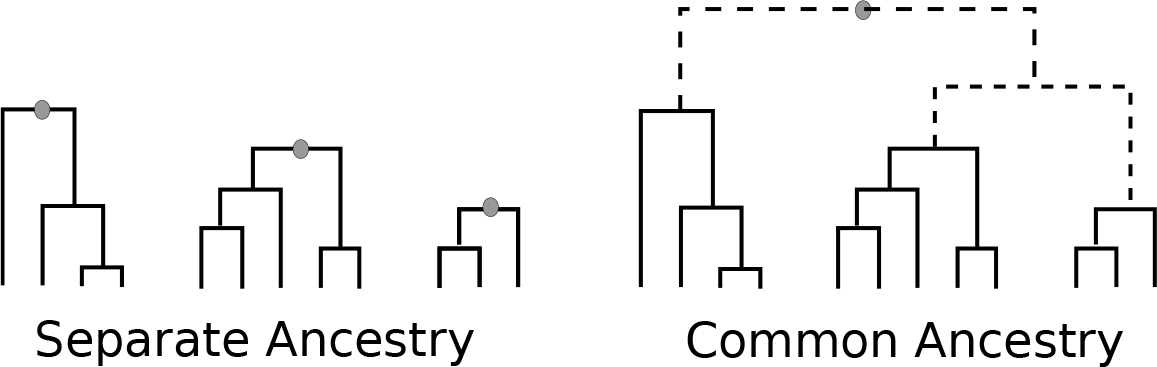
Common and Separate Ancestry Models. In the separate ancestry likelihood model (left), ancestral sequences (gray circles) are unrelated subject to constraints within each group. In the common ancestry model (right), there is a single ancestral sequence. Dashed branches represent the tree that connects the ancestors of each group.

The BIC method estimates a single tree topology from all 178 taxa using maximum likelihood on the concatenated data set. Within this tree, each primate suborder and family is monophyletic. The maximum likelihood tree topologies for any group of taxa in both the CA and SA models are assumed to match the corresponding subtree from this single maximum likelihood tree, but branch length parameters are estimated separately for each gene and the subset of taxa with data for that gene. The CA model includes a single nucleotide substitution model for each gene. The SA models include separate substitution models for each gene and group. Both the SA and CA models have a separate stationary distribution for each site. In the CA model, there is a single ancestral sequence, represented graphically as a single gray dot on the right tree in Figure 1, drawn at random from the empirical distributions by site across all taxa. In the SA model, there are separate draws from the empirical distributions by site across all taxa for the ancestor of each group, represented by the multiple gray dots on the left trees in Figure 1. For both models, the likelihood is recalculated for each site. The BIC score is computed for each model as

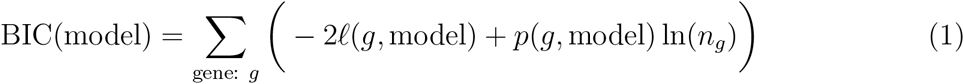

where ℓ(g, model) is the (natural) log likelihood for the corresponding model and gene *g, p*(*g*, model) is the number of parameters estimated for gene *g* for the given model, and *n_g_* is the number of sites for gene *g*. The model with the lowest BIC score has the most support and the difference in BIC scores between models determines the posterior probability of each model given equal prior probabilities of each. Further details are in Methods.

## Parsimony difference test for primate common ancestry

The parsimony difference test uses as a test statistic the portion of the parsimony score that can be attributed to changes among the group ancestors and not within the groups. We calculate this by subtracting the sum of parsimony scores maximized over individual groups from the maximum parsimony score of the best tree for all taxa. The parsimony difference score may be thought of as the total number of nucleotide substitutions that would be mapped to the best tree relating group ancestors (the dashed branch tree in the right of Figure 1). As universal CA posits that group ancestors are related by such a tree and SA does not, CA predicts this score will be smaller than SA does.

In order to test SA versus CA, we treat SA as the null hypothesis and CA as the alternative. Consequently, all calculations of p-values assume SA. In brief, separately for each SA group we fit a maximum likelihood model of nucleotide substitution that accomodates site-specific information of nucleotide base usage to account for functional and biological constraints. We then simulate a large number of data sets for each group under these fitted models and approximate via this simulation the null sampling distribution of the test statistic. This allows us to assess significance of the actual data relative to these hypotheses. Model and calculation details are in Methods.

We examined common ancestry among primates at three different taxonomic levels. First, we examined the base of the primate tree and asked if strepsirrhines and haplorrhines have a common ancestor. Second, we looked at an intermediate (family) level of taxonomy and considered the possibility of SA among the 16 primate families. Finally, we examined the special case of *Homo sapiens* and tested if separate creation of humans can be rejected to the alternative that humans and all other primates share a common ancestor.

## RESULTS AND DISCUSSION

### BIC test of common ancestry

Table 2 contains a summary of the BIC scores for the CA model, the family SA model, and the suborder SA model. BIC scores are calculated for each model according to Equation 1. Each BIC score is a sum over BIC scores from each of the 54 genes. A table with the log-likelihoods and parameter counts for each gene is in supplementary material. The CA model provides the best explanation of the data among the models explored. In a direct comparison between the CA model and the family SA model, an observer that thought *a priori* that both models were equally valid will, after evaluating the sequence data, be essentially certain that CA is correct: the posterior probability for the family SA model is about e^−20,854.95/2^ = 10^−4528.6^. Similarly, CA is essentially certain relative to suborder SA which has posterior probability about 10^−124.6^.

**Table 2:**
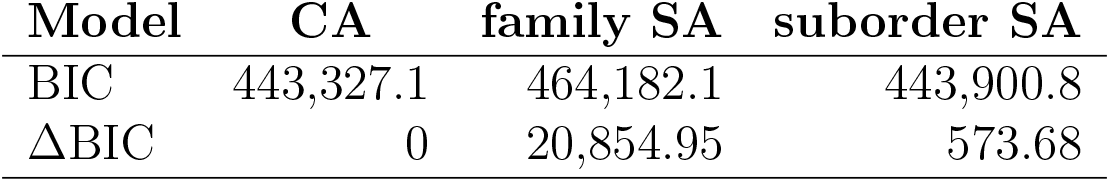
Results of the BIC Test. Each of the three models uses the same empirical relative frequency of called bases at each site across all primates as the maximum likelihood estimate for the site-specific nucleotide base probabilities. Each model estimates branch lengths and nucleotide substitution parameters separately for each of the 54 genes. The CA model assumes a single tree and common ancestry for all taxa. The family SA model assumes independent origins for each of the 16 primate families. The suborder SA model assumes separate origins for both primate suborders.

| Model | CA | family SA | suborder SA |
| --- | --- | --- | --- |
| BIC | 443,327.1 | 464,182.1 | 443,900.8 |
| ΔBIC | 0 | 20,854.95 | 573.68 |

### Parsimony Difference Test: Strepsirrhini/Haplorrhini Common Ancestry

The maximum likelihood trees for the 140 taxa in the haplorrhini and 38 strepsirrhini suborders, pruned to the family level for simplicity, are displayed in Figure 2. Files containing the full trees with estimated branch lengths, estimated nucleotide substitution parameters, and estimated site-specific ancestral sequence probability distributions are in supplementary materials. The null sampling distribution of the parsimony difference test statistic, as determined by a parametric bootstrap sample of 1000 simulated data sets, is shown in Figure 3 and has mean 3415.0 and standard deviation 40.0. The corresponding test statistic is 1097 which is 57.9 standard deviations below the mean. Assuming the left tail of the null sampling distribution is described well by a normal distribution, the corresponding p-value is about 10^-1680^. There is overwhelming support against separate ancestry in favor of common ancestry between the two primate orders. To consider this evidence from another perspective, if haplorrhini and strepsirrhini had unrelated ancestors whose ancestral sequences were constrained in similar fashion to their modern day descendendents, then in the context of a best-fitting likelihood model of nucleotide substitution, we would expect the unobserved ancestral sequences to differ in about 3500 sites (out of the nearly 35,000 sites considered), give or take a hundred or so. However, the actual data is consistent with ancestral sequences that differ in only about 1100 sites, which is much more plausibly explained by descent from a common ancestor than by chance. Indeed, the probability of the observed result under our SA model is about the same as that of correctly choosing at random one atom from all of the approximate 10^80^ atoms in the visible universe 21 times in a row.

**Figure 2:**
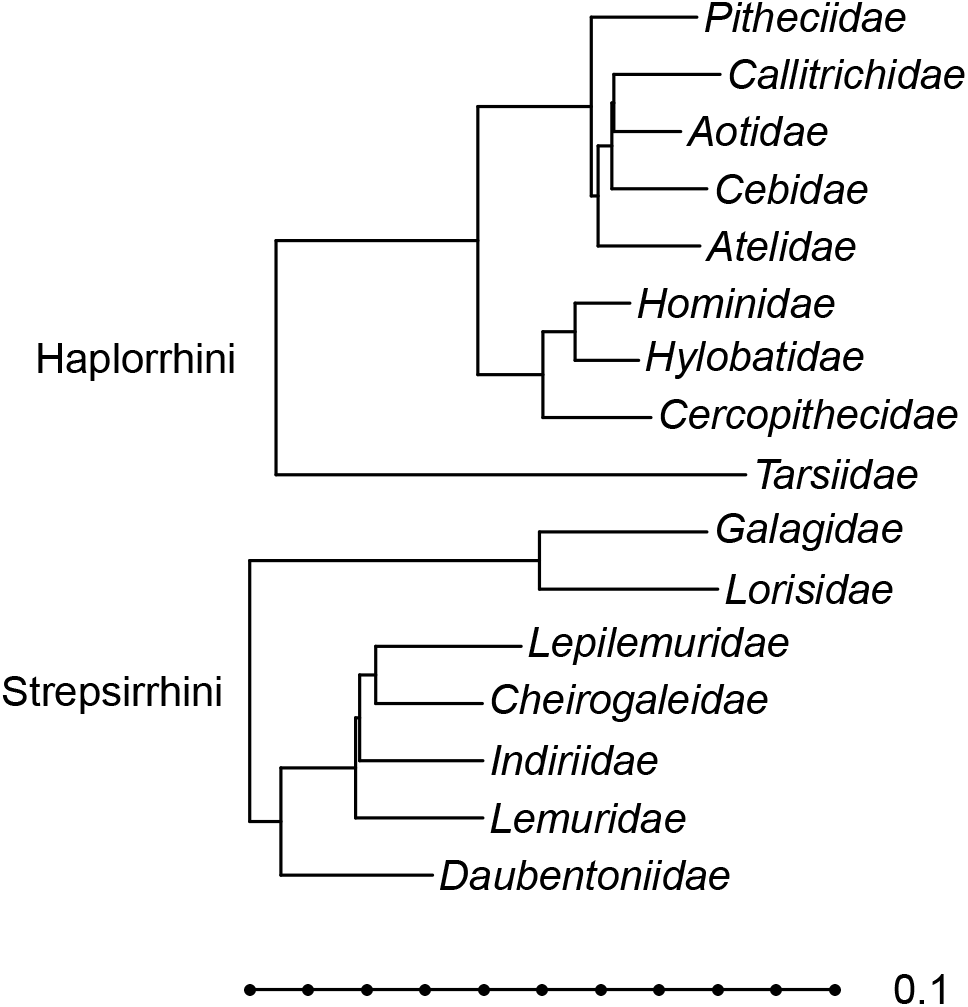
Haplorrhini and Strepsirrhini Trees. The maximum likelihood trees for each primate order, pruned to families, are displayed on the same scale. The entire scale bar has length 0.1 expected substitutions per site.

**Figure 3:**
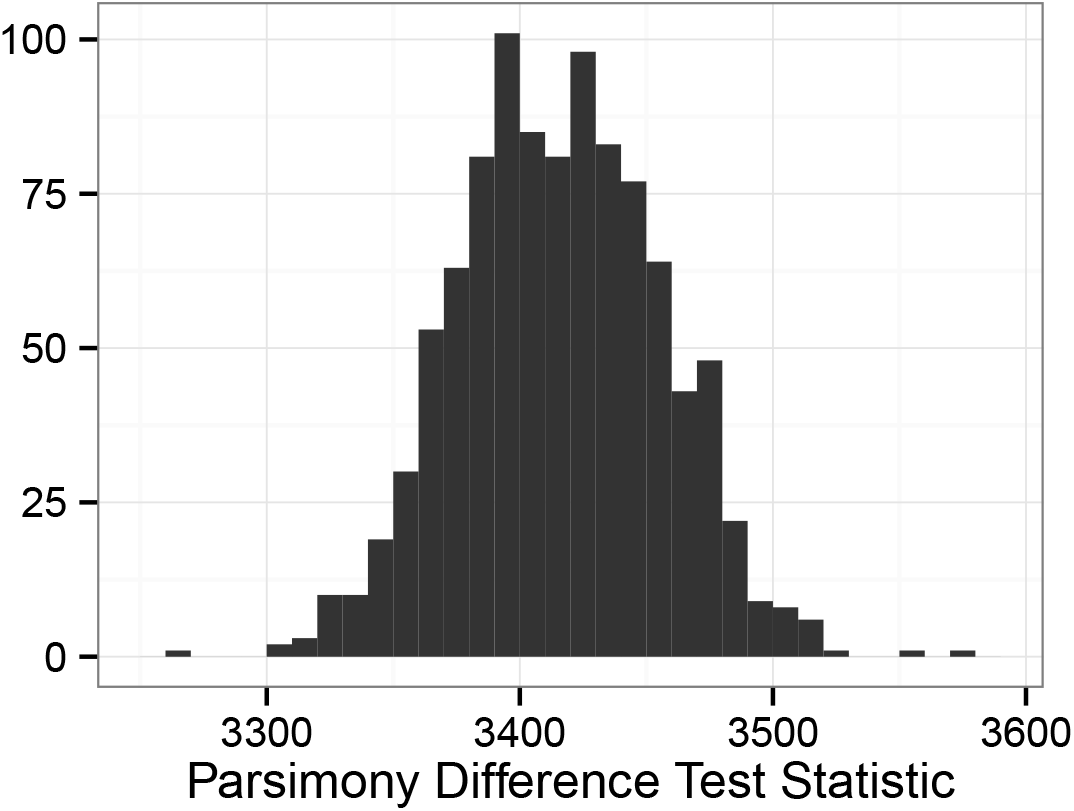
Haplorrhini/Strepsirrhini separate ancestry null distribution. The sampling distribution of the parsimony difference test statistic in the test of separate ancestry in the primate orders Happlorrhini and Strepsirrhini based on a parametric bootstrap sample of 1000 has mean 3415.0 and standard deviation 40.0.

### Parsimony Difference Test: Primate Family Common Ancestry

We next test SA among the 16 primate families from Table 1. Maximum likelihood estimates of primate family trees and branch lengths and family-level estimates of nucleotide substitution model parameter values and site-specific ancestral sequence nucleotide base probability distributions are included in supplementary materials. The null sampling distribution of the parsimony difference test statistic is shown in Figure 4 and has mean and standard deviation 13,422.3 and 45.9, respectively. The actual data has a parsimony difference score of 10,125, which is 71.8 standard deviations below the mean. Assuming accurate normal behavior in the tail of the true sampling distribution corresponds to a p-value of 10^-2581^. The evidence against SA among primate families is even stronger than the evidence against SA between the two primate orders. SA predicts that the best possible tree connecting primate family ancestral sequences would contain well over 13,300 sequence changes on the tree. However, the real data can explain differences among these ancestral sequences with only just over 10,000 changes. Acceptance of SA means believing a chance occurrence less likely than correctly identifying an atom from the universe 32 times in succession has occurred, which is truly overwhelming evidence against SA in favor of CA among primate families.

**Figure 4:**
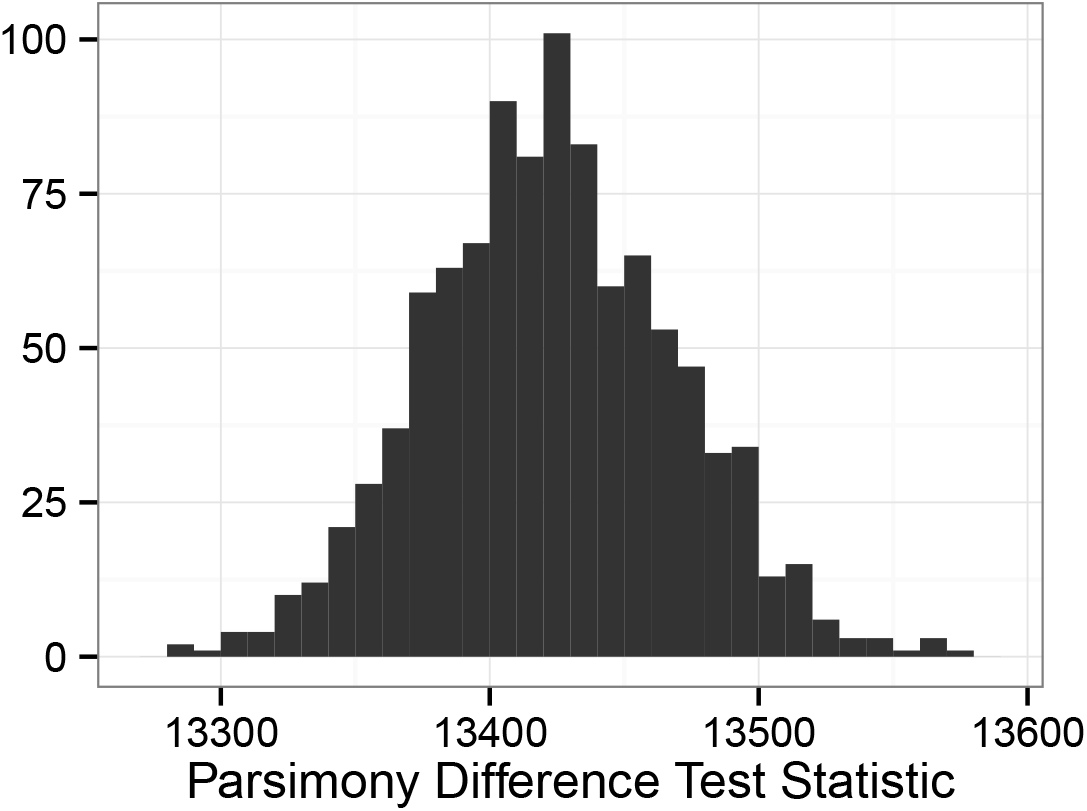
Primate family separate ancestry null distribution. The sampling distribution of the parsimony difference test statistic in the test of separate ancestry among 16 primate families based on a parametric bootstrap sample of 1000 has mean 13,422.3 and standard deviation 45.9.

### Parsimony Difference Test: Human Common Ancestry

Finally, we turn to the question of SA for *Homo sapiens* from all other primates. For this test we consider a maximum likelihood tree for the 177 non-human primates in this data set. Summaries of this tree and associated substitution model parameters and ancestral sequence distribution probabilities are in supplementary materials. The one-taxon human maximum likelihood tree is a single node with no branches where the human sequence is identical to its ancestor sequence. Effectively, the method of this paper for this case involves simulating sequences for all non-human primates and determining the parsimony cost of adding the genuine human sequence to the best tree among other primates for each simulated data set.

The sampling distribution of the test statistic has mean 2203.3 and standard deviation 22.0 (see Figure 5). The real human sequence adds only 136 to the parsimony score when aligned with the real primate sequence data, which is 93.9 standard deviations below the mean. The normal distribution approximation for the p-value is 10^-4413^ and the evidence against SA of humans and other primates is overwhelming. The chance of simulating data in the nonhuman primate tree as close to the real human sequence as the actual primate data is uncomprehendlingly small. While accounting for site-by-site sequence constraints among primates, SA predicts that the human sequence will differ by about 2200 changes from the best place it might attach to a simulated primate tree. The actual small distance between the real human sequence and the real sequences of other primates is wholly inconsistent with the assumption of SA, and is explained most plausibly by CA.

**Figure 5:**
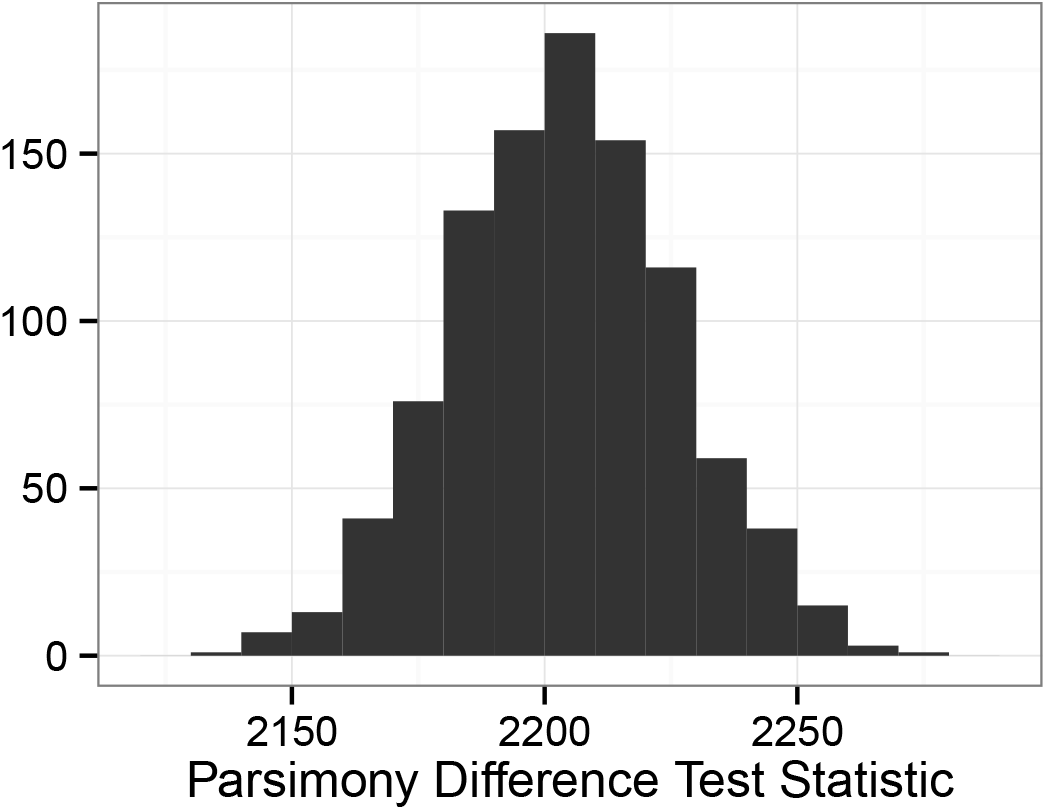
Human separate ancestry null distribution. The sampling distribution of the parsimony difference test statistic in the test of separate ancestry for *Homo sapiens* based on a parametric bootstrap sample of 1000 has mean 2203.3 and standard deviation 22.0.

We considered one additional alternative test of human SA. For this test, we simulated 1000 pseudo-human sequences by replacing non-gap human sequence nucleotides with called bases selected at random from the other primate sequences at each site, creating a collection of chimeric primate sequences. For each such sequence, we found the parsimony difference statistic with the original non-human primates and examined their distribution. The mean and standard deviation were XXX and XXX. The real human sequence score is XXX standard deviations below the mean. We infer from this test that the real human sequence is better explained by CA with all primates, because sequences generated independently are much further from other primate sequences than is the genuine human sequence, even while adjusting for site-specific nucleotide base usage patterns.

## CONCLUSIONS

We have developed novel statistical approaches to test CA versus SA from aligned DNA sequences based on maximum likelihood estimation, BIC, and parametric bootstrapping of a parsimony difference test statistic. Our model treats nucleotide base probabilities separately at each site in order to account for biological constraints that limit nucleotide usage differently by site.

We find overwhelmingly strong evidence against SA in favor of CA in primates at both the subordinal and family levels. Additionally, we find common ancestry between primate orders and among primate families. We find very strong statistical evidence against a hypothesis of SA of humans from other primates, This supports the conventional view that humans are closely related to other primates rather than deriving from an independent origin event.

## METHODS

### Data Preparation

We obtained an alignment that consists of 54 nuclear genes and 191 taxa including 186 primate taxa for a total of 34,941 base pairs (*Perelman et al.* 2011). We modified the alignment to meet our needs to examine CA among primates first by eliminating the five outgroup species. Then, we eliminated the 81 sites with no called bases — these were sites where the outgroups had sequence data, but the primates did not. Next, to eliminate potential bias in our tests due to the inclusion of multiple subspecies of a single species, we created a single sequence per species in such a way as to maximize retained information. This process entailed combining six pairs of taxa and one triplet of taxa, reducing the number of taxa in our study by eight. Specifically, we combined sequences from: (1) *Pan troglodytes verus* and *Pan troglodytes troglodytes*; (2) *Aotus azarae, Aotus azarae boliviensis,* and *Aotus azarae infulatus*; (3) *Eulemur macaco* and *Eulemur macaco flavifrons*; (4) *Microcebus murinus ssp. 1* and *Microcebus murinus ssp. 2*; (5) *Propithecus verreauxi* and *Propithecus verreauxi coquereli;* (6) *Semnopithecus entellus* and *Semnopithecus entellus entellus;* (7) *Varecia variegata rubra* and *Varecia variegata variegata.* Sequence regions from subspecies where both (or all three) are determined, agree for about 99.9 percent of sites, so we are confident that using sequence data from one subspecies where another is missing provides a highly accurate reconstruction. In the few sites where measured bases disagreed, we used the more specific base when the base codes were compatible (for example, A and R became A) and introduced base uncertainty when there was discordant information (for example, C and T became Y). The final alignment contained 178 species and 34,860 sites.

### Models for the BIC test

The BIC test requires likelihood models for the CA and both SA hypotheses. We found the maximum likelihood tree using the full concatenated aligned data set using RAxML version 8.0.24 (Stamatakis 2014). Each suborder and each family from Table 1 was monophyloletic in this tree topology, which we used as a constraint for subsequent analyses. We estimated branch lengths and substitution model parameters separately for each gene and hypothesis using RAxML and the GTR + Γ model of nucleotide substitution in the following way. For the CA model, we estimated branch lengths for a single tree for all available taxa and parameters from a single substitution model. The suborder and family SA models estimated separate branch length and substitution model parameters for each group (2 groups and up to 16 groups, respectively). One exception was for families with fewer than four taxa with data available for a given gene, which was too few taxa to run RAxML. In these cases, branch lengths and substitution parameters were used from the entire concatentation of data for two or three taxa. With zero or one taxa, there are no branch lengths or substitution parameters to estimate. Another exception was when one or more GTR parameters were estimated as zero, which happened for some short genes and small families. In this case, we replaced parameter values with GTR parameter estimates using all data from the gene. We then recalculated the likelihood of each tree for each model and gene with these estimated branch lengths and substitution model parameters, but replaced each stationary distribution with the empirical relative frequencies at each site for each model and group, using the **pml()** function in R package **phangorn** (Schliep 2011; R Core Team 2013).

For a gene with m total taxa and n sites, the CA, suborder SA, and family SA models all have 3*n* parameters for the site-specific stationary distributions. The CA model also has 2*m* – 3 branch length parameters and 6 parameters for the substitution model (less the stationary distribution) for a total of 2*m* + 3*n* + 3. The suborder SA model has 2*m* – 6 branch lengths and 12 substitution model parameters for a total of 2*m* + 3*n* + 6. For a gene with sequences from *x* families, the family SA model has 2*m* – 3*x* branch lengths and 6*x* substitution model parameters, totalling 2*m* + 3*n* + 3*x* parameters before reducing for the exceptions mentioned earlier. Using the likelihoods and parameter counts for each gene and model, the BIC score for each model is calculated using Equation 1.

### Models for the parsimony difference test

The CA hypothesis assumes that group ancestors descended in a tree-like fashion from a single common ancestor and that all species are related by a single universal tree. The probability distribution of the one ancestral sequence at the root of the tree is independent for each site, but not identically distributed. Each site has a separate probability distribution for the ancestral state that is estimated as the conditional distribution of the ancestral state given the maximum likelihood tree and the observed sequence data at each site. Sequences at group ancestors are modeled to have undergone the same substitution process as within groups while evolving along a tree from a single common ancestor.

The SA hypothesis assumes that the group ancestral sequences are created separately and independently. The distribution of each separate ancestral sequence is estimated on a site-by-site basis as the conditional distribution of the ancestral state given the maximum likelihood tree for the group and the observed sequence data within the group. Furthermore, the SA hypothesis assumes separate parameters for the nucleotide substitution process for each group. We assume a null hypothesis of SA and determine if there is sufficient evidence in the data to reject SA in favor of CA.

Sequence changes within groups are modeled to follow a conventional phylogenetic likelihood model of nucleotide substitution, the general time reversible model with gamma-distributed rate variation among sites (GTR+Γ). We used RAxML version 8.1.17 (Stamatakis 2014) to estimate phylogenies and nucleotide substitution parameters. Unlike conventional likelihood models, we use the existing sequence data to estimate the ancestral sequence for simulation instead of sampling the ancestral sequence from the stationary distribution. We do this so that nucleotide base usage in simulated sequences mimics the actual usage on a site-by-site basis in order to better simulate realistic sequences that are more likely to adhere to unspecified biological and functional constraints.

Estimated trees are unrooted. We root the trees using midpoint rooting in which the midpoint is midway on the longest path between any two tips. Given a rooted estimated likelihood tree, estimated substitution model parameters, and an alignment, for each site we calculate separately the conditional distribution of the nucleotide base at the ancestral root. The algorithm for this calculation is similar to the first step of simulating unobserved sequences in (Nielsen 2002). As the total distance from ancestral tree root to tips of the tree is fairly short (much less than one expected change per site), the effect is that while simulating sites where observed data show no variation, the simulated ancestral state is overwhelmingly likely to agree with the observed data and there is a very high chance that the simulated bases at the tree tips will also show no variation.

The real data also has characteristics of missing data. As not all genes were sequenced for all taxa and as a history of small insertions and deletions require the addition of gaps in the alignment, the real alignment has symbols ‘-’ to represent insertions or deletions and ‘?’ to represent nucleotides assumed to be present, but unmeasured. Models for the substitution process only, as in this paper, treat these forms of missing data identically. Furthermore, a small fraction of the alignment contains symbols such as ‘R’ or ‘Y’ to indicate uncertainty in the called base. We do not attempt to simulate insertions and deletions, but rather overwrite in simulated data gaps and question marks where the real data has such missing data. We use partially called bases when estimating ancestor sequence probabilities, but replace all such symbols with gaps, in both simulated and real alignments, before computing parsimony scores to eliminatate the potential effects of such partially called bases.

To illustrate how features in the real data are preserved in the simulations, the following shows the first 11 sites of the *ABCA1* gene for the simulated *Homo sapiens* sequence from the parsimony difference test of SA between primate orders, which includes a gap in the 5th site, and corresponds to the real human sequence aligned as TGTG-TCTCAC.

1. TCTG-TCTCAA
2. GGTG-TCTCAC
3. TGTG-CCTCAA
4. TGTG-TCTCAC
5. CATG-CCTCAG
6. GATG-TCTCAG
7. TATG-TCTCAA
8. TGTG-TCTCGG
9. TATG-TCTCAA
10. CGTG-CCTCAC

The simulated haplorrhini ancestor sequence at these sites is drawn independently at each site from the following distributions.

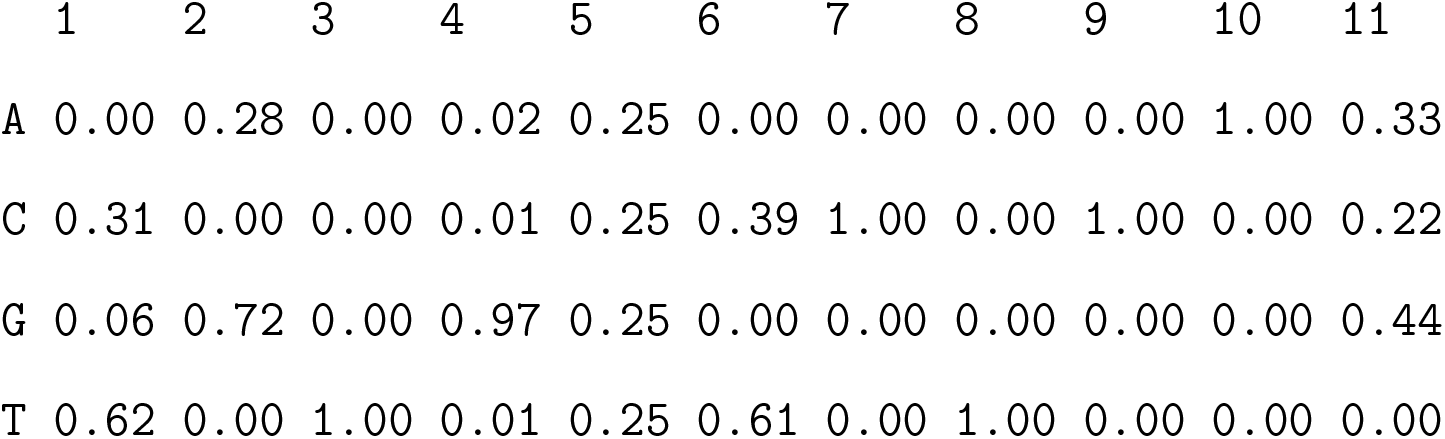

We see that the simulated sequences tend to contain nucleotides at each site that are common among the genuine haplorrhini sequences. For example, at sites 3 and 7-10, all haplorrhini have the same nucleotide and the simulated human sequence almost always agrees. There is one exception in site 10 of the 8th simulated sequence. Here, even though the simulated ancestor sequence was certainly an A, the simulated rate for this site and simulation must have been large enough for a substitution to occur. Similarly, there will be cases where the simulation at a site has no variation, but the real data does vary, if the simulated substitution rate is very small. But for the most part, simulated human sequences have similar base usage to the observed haplorrhini sequences at each site and simulated sites mimic the genuine sequences.

### Testing Separate Ancestry

To test SA, we do the following:

1. Partition the taxa into groups for which SA is to be tested.
2. Find maximum likelihood tree and substitution model parameter values separately for each group.
3. Root each tree by midpoint rooting.
4. Determine the ancestral sequence probability distributions for each group and each site.
5. Simulate an ancestral sequence for each group.
6. Use the estimated substitution model to simulate sequences to the tips for each group's tree.
7. Replace nucleotides with gaps in the the simulated alignments where the real alignment has gaps or uncertainty for the taxon in question.
8. Calculate parsimony scores (as described below) for each group alignment separately and for the single combined alignment across all groups.
9. Compute the parsimony difference score: the parsimony score of the combined alignment minus the sum of the parsimony scores of all groups.
10. Repeat steps 5-9 1000 times.
11. Add gaps for partially resolved bases in the original alignment and compute parsimony difference statistic from the original data.
12. Determine z-score of the observed test statistic and find approximate normal-based p-value.

We used PAUP* version 4.0b10 for Unix (Swofford 2003) to do all parsimony calculations. Sequence simulations, graphs, and p-value calculations use R version 3.02 (R Core Team 2013) and the **ape** package for phylogenetics (*Paradis et al.* 2004). Simulating new data from a fitted maximum likelihood model for testing is an application of the parametric bootstrap (Efron and Tibshirani 1993).

## References

Baum, D. A., C. Ané, B. Larget, C. SolíS-Lemus, L. S. T. Ho, P. Boone, C. Drummond, M. Bontrager, S. Hunter, and B. Saucier. 2015. Statistical evidence for common ancestry: Application to primates. XXX XX:X–XX.

Bontrager, M., B. Larget, C. Ané, and D. A. Baum. 2015. Statistical evidence for common ancestry: Testing for signal in silent sites. XXX XX:X–XX.

Coyne, J. A. 2009. Why Evolution is True. Viking Penguin, New York.

Darwin, C. 1859. On the Origin of Species by Means of Natural Selection, or the Preservation of Favoured Races in the Struggle for Life. John Murray, London.

De Oliveira Martins, L. and D. Posada. 2014. Testing for universal common ancestry. Systematic Biology 63:838–842. DOI:10.1093/sysbio/syu041.

Efron, B. and R. J. Tibshirani. 1993. An Introduction to the Bootstrap, volume 57 of Monographs on Statistics and Applied Probability, chapter 16. Chapman and Hall, New York.

Eugene V. Koonin, Y. I. W. 2010. The common ancestry of life. Biology Direct 5. Http://www.biology-direct.com/content/5/1/64.

Groves, C. P. 1991. A Theory of Human and Primate Evolution. Clarendon Press, Oxford.

Hill, W. C. O. 1966. Primates Comparative Anatomy and Taxonomy VI Catarrhini Cercopithecoidea Cercopithecinae. Edinburgh University Press, Edinburgh.

Nielsen, R. 2002. Mapping mutations on phylogenies. Systematic Biology 51:729–739.

Paradis, E., J. Claude, and K. Strimmer. 2004. APE: analyses of phylogenetics and evolution in R language. Bioinformatics 20:289–290.

Penny, D., L. R. Foulds, and M. D. Hendy. 1982. Testing the theory of evolution by comparing phylogenetic trees constructed from five different protein sequences. Nature 297:197–200.

Penny, D., M. D. Hendy, and A. M. Poole. 2003. Testing fundamental evolutionary hypotheses. Journal of Theoretical Biology 223:377–385.

Perelman, P., W. E. Johnson, C. Roos, H. N. Seuénez, J. E. Horvath, M. A. M. Moreira, B. Kessing, J. Pontius, M. Roelke, Y. Rumpler, M. P. C. Schneider, A. Silva, S. J. Obrien, and J. Pecon-Slattery. 2011. A molecular phylogeny of living primates. PLoS Genetics 7. E1001342.

R Core Team. 2013. R: A Language and Environment for Statistical Computing. R Foundation for Statistical Computing, Vienna, Austria.

Schliep, K. P. 2011. phangorn: phylogenetic analysis in R. Bioinformatics 27:592–593.

Sober, E. and M. Steel. 2002. Testing the hypothesis of common ancestry. Journal of Theoretical Biology 218:395–408.

Stamatakis, A. 2014. RAxML version 8: a tool for phylogenetic analysis and post-analysis of large phylogenies. Bioinformatics 30:1312–1313.

Swofford, D. L. 2003. PAUP*. Phylogenetic Analysis Using Parsimony (*and Other Methods). Version 4. Sinauer Associates, Sunderland, Massachusetts.

Theobald, D. L. 2010a. A formal test of the theory of universal common ancestry. Nature 465:219–222. Doi:10.1038/nature09014.

Theobald, D. L. 2010b. Theobald reply. Nature 468. Article ID E10.

W. Timothy J. White, D. P., Bojian Zhong. 2013. Beyond reasonable doubt: Evolution from DNA sequences. PLoS ONE 8. E69924.

Yonezawa, T. and M. Hasegawa. 2010. Was the universal common ancestry proved? Nature 468. Article ID E9.

Yonezawa, T. and M. Hasegawa. 2012. Some problems in proving the existence of the universal common ancestor of life on earth. The Scientific World Journal 2012. Article ID 479824, 5 pages doi:10.1100/2012/479824.

